# Ectopic cervical thymi and no thymic involution until midlife in naked mole-rats

**DOI:** 10.1101/2021.08.01.454657

**Authors:** Stephan Emmrich, Frances Tolibzoda Zakusilo, Alexandre Trapp, Xuming Zhou, Quanwei Zhang, Ellen M. Irving, Michael G. Drage, Zhengdong Zhang, Vadim N. Gladyshev, Andrei Seluanov, Vera Gorbunova

**Affiliations:** Department of Biology, University of Rochester, Rochester NY 14627, USA; CAS Key Laboratory of Animal Ecology and Conservation Biology, Institute of Zoology, Chinese Academy of Sciences, Beijing 100101, China; Department of Genetics, Albert Einstein College of Medicine, New York City, NY, USA; Pathology and Laboratory Medicine, University of Rochester Medical Center, Rochester, NY, USA; Division of Genetics, Department of Medicine, Brigham and Women’s Hospital, Harvard Medical School, Boston, MA, USA

**Keywords:** Naked Mole-Rat, Aging, Immunosenescence, Lymphopoiesis, Thymus, T-Lymphocytes, CD4- CD8 Ratio

## Abstract

Immunosenescence is a hallmark of aging and manifests as increased susceptibility to infection, autoimmunity, and cancer in the elderly. One component of immunosenescence is thymic involution, age-associated shrinkage of the thymus, observed in all vertebrates studied to date. The naked mole-rat (*Heterocephalus glaber*) has become an attractive animal model in aging research due to its extreme longevity and resistance to disease. Here we show that naked mole rats display no thymic involution up to 11 years of age. Furthermore, we found large ectopic cervical thymi in addition to the canonical thoracic thymus, both being identical in their cell composition. The developmental landscape in naked mole-rat thymi revealed overt differences from the murine T cell compartment, most notably a decrease of CD4^+^/CD8^+^ double-positive cells and lower abundance of cytotoxic effector T cells. Our observations suggest that naked mole rats display a delayed immunosenescence. Therapeutic interventions aimed at reversing thymic aging remain limited, underscoring the importance of understanding the cellular and molecular mechanisms behind a sustained immune function in the naked mole rat.

## Introduction

Age-related loss of cellularity and weight of the thymus, termed thymic involution, occurs in all mammals (Shanley et al., 2009). Thymic involution contributes to immunosenescence, leading to increased incidence of cancers, autoimmunity, and opportunistic infections in the elderly (Dixit, 2012, Lynch et al., 2009). Thymic involution remains an evolutionary mystery since it occurs in most vertebrates despite its negative effects. In vertebrates aging of the thymus gland precedes age-associated phenotypic changes in other organs, with thymic function gradually decreasing from the first year of human life (Shanley et al., 2009). The process of human thymic involution is incompletely understood, in part due to a lack of animal models showing delayed immunosenescence (Tong et al., 2020).

Naked mole-rats (NMR) display exceptional longevity, with a maximum lifespan of 32 years (Buffenstein, 2008, Buffenstein & Jarvis, 2002), without increased mortality due to aging (Ruby et al., 2018). In comparison, a similarly sized house mouse has a maximum lifespan of 4 years (de Magalhaes et al., 2005, Turturro et al., 1999). NMRs display negligible senescence (Buffenstein, 2008), characterized by very slow changes in physiological parameters with age, and the lack of an age-related increase in mortality rate (Finch, 1990). Accordingly, NMRs do not exhibit age related changes in basal metabolism, body composition, or bone mineral density (O’Connor et al., 2002, Buffenstein & Ruby, 2021). NMRs show a very low incidence of cancer (Buffenstein, 2008, Delaney et al., 2013, Delaney et al., 2016). NMRs produce abundant high molecular weight hyaluronic acid (HMW-HA) responsible for their resistance to solid tumors (Tian et al., 2013a, Tian et al., 2015) and feature a serum metabolome resembling calorically restricted mice(Lewis et al., 2018, Puppione et al., 2021). Thus, NMRs present a unique model to study the mechanisms of healthy longevity (Edrey et al., 2011, Gorbunova et al., 2014). However, the information about the immune system of NMRs is limited. Very recently, whole spleen single cell RNA-Sequencing revealed the absence of canonical Natural Killer Cells (NKC) in NMRs (Hilton et al., 2019). It was also reported that blind mole-rats, an unrelated clade of subterranean rodents that convergently evolved extreme longevity, sustain sustained T cell repertoire diversity in old age (*Spalax spp.*)(Izraelson et al., 2018). At present there is, however, no data regarding the T cell compartment in NMRs.

Here we set out to characterize the thymus and T-Lymphopoiesis in long-lived NMRs in comparison with short-lived mice. Our results revealed unexpected presence of additional cervical thymi in the NMR, and the absence of thymic involution until 11 years of age, which is the oldest age of animals in our research colony.

## Results

Thymic involution and decline of T cell function predisposes to opportunistic diseases and presents a major risk factor in the elderly (Dixit, 2012, Torroba & Zapata, 2003). Since NMRs are characterized by negligible senescence, we set out to examine whether they display thymic involution. To enable the analysis of the NMR hematopoietic cells, we previously developed a flow cytometry (FACS) staining with cross-reactive monoclonal antibodies (moAbs) and functionally characterized purified hematopoietic stem and progenitor (HSPC) fractions (Emmrich et al., 2019). Through CITE-Seq we showed that naked mole-rat T cells (TC) express CD3, LAT and LCK are immunophenotypically Thy1.1^int^/CD34^−^/CD11b^−^ (**Figure S1a, b**).

Thymus glands extracted from the thoracic cavity of young NMRs and mice both featured similar sized lymphocytes and largely absent myeloid cell fractions (**Figure S2a**). When we extracted lymph nodes (LN) from 43 naked mole-rat necks we noted two types of cervical nodes (**Figure 1a**): the first type corresponding to LN, and the second resembling a thymus, which we termed cervical thymi. Thoracic and cervical thymi contained a CD34^+^ fraction while LNs did not, and shared histological features such as many medullary sinuses, no germinal centers and more basophilic cytoplasms than LNs (**Figure 1b-c**). Conceivably, both thymi types stain positive for cytokeratin as seen for mouse and human thymi, while LNs are cytokeratin^−^ with a much lower TC:BC ratio (**Figure 1d, Figure S2b**). Embryonic thymus development proceeds through detachment of the primordium, which contains thymus- and parathyroid-anlagen, from the 3rd pharyngeal pouches, followed by separation of the anlagen and migration of thymic lobes into the chest cavity (Gordon & Manley, 2011). We compared the macroscopic anatomy with mice and found undersized thoracic thymus lobes in NMR neonates (**Supplementary S2c**). To our surprise, the cervical thymus was clearly visible in the ventral pharyngeal region, and appeared markedly larger than their thoracic thymus (**Figure S2d**).

**Figure 1.**
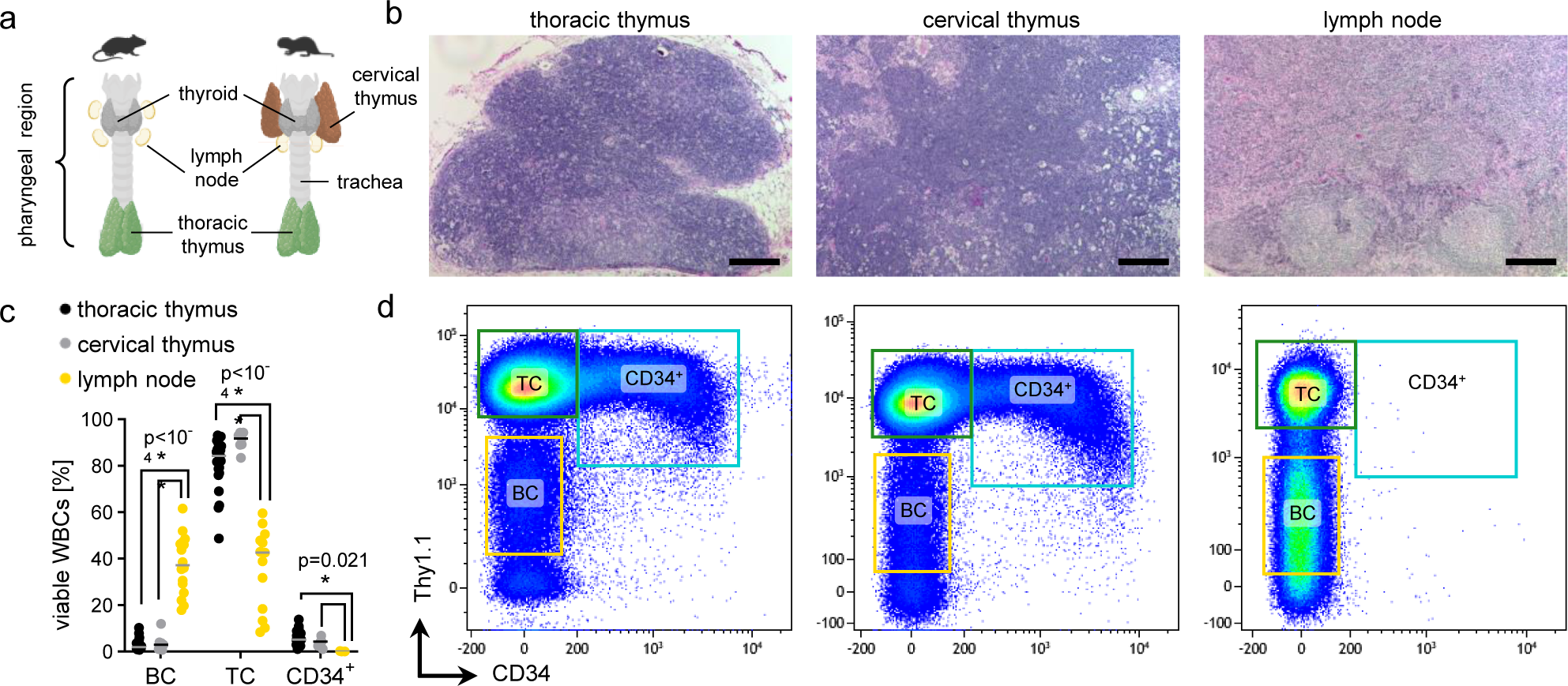
Naked mole-rats have an additional thymus. **a**, Schematic of the throat anatomy between mouse and naked mole-rat, highlighting lymphoid organs. **b**, Hematoxylin & Eosin stained tissue sections of naked mole-rat thoracic [left] and cervical [middle] thymus or lymph node [right] (LN); Scale bars 200µm. **c**, Frequencies of cell populations across lymphatic organs. Thoracic, n=31; cervical, n=16; B lymph node, n=7. **d**, FACS gating of viable leukocytes of naked mole-rat thoracic [left] and cervical [middle] thymus or LN [right].

### Conserved thymopoiesis between mouse and naked mole-rat

Next, we performed whole thymus CITE-Seq from two 3 month-old and two 12 month-old mice versus two 3 year-old and two 11 year-old NMRs (**Figure S3a**). The classical thymus-specific FACS pattern of CD4^+^/CD8^+^ double-positive (DP) thymocytes in mice (**Figure S3b**) was resolved into 13 cell states of 20019 single cell transcriptomes (**Figure 2a**). We captured sorted naked mole-rat CD34^+^ thoracic thymus cells (**Figure S3c**) and two LNs from the younger animals and integrated those with naked mole-rat thoracic and cervical thymi. The integrated dataset comprised 70928 cells across 11 droplet libraries from 3 tissues and one FACS-purified sample (**Figure 2b**). We found 15 distinct communities which were annotated based on canonical marker gene expression for respective TC subsets (**Figure 2c**). The earliest developmental stage in the thymus are CD4^−^/CD8^−^ double-negative (DN) early T cell progenitors (ETP), while the expression of CD44 and CD25 further subdivides the DN state. A DN2/3 cluster was the earliest distinct developmental time point detected, in mice showing exclusive CD25^hi^ expression, a Kit high-to-low-expression gradient and a fraction of CD44^lo^ cells as measured by CITE-signals (**Figure S3d**). In both species expression of PTCRA, NOTCH1 and its target HES1 was conserved in DN2/3 cells (**Figure S3e-f, Supplementary Table 1**). The transient DN4 population (CD44^−^/CD25^−^) initiates TCR-α gene rearrangements and upregulates expression of CD4 and CD8 to yield DPs, which usually progress through an immature cycling CD8^+^ intermediate single-positive population (ISP)(MacDonald et al., 1988). We found an equivalent cell state DN4/ISP in both species, which was marked by overexpression of cell cycle genes, retention of PTCRA and rising expression of CD4/CD8. Remarkably, sorted CD34^+^ naked mole-rat thymocytes were strongly enriched for DN2/3 and DN4/ISP clusters (21%, 35%; **Figure S3g**), while CD34 mRNA and CITE-signal were specific to DN2/3. This trait is shared with humans, who maintain CD34^+^ primitive ETPs (Terstappen et al., 1992).

**Figure 2.**
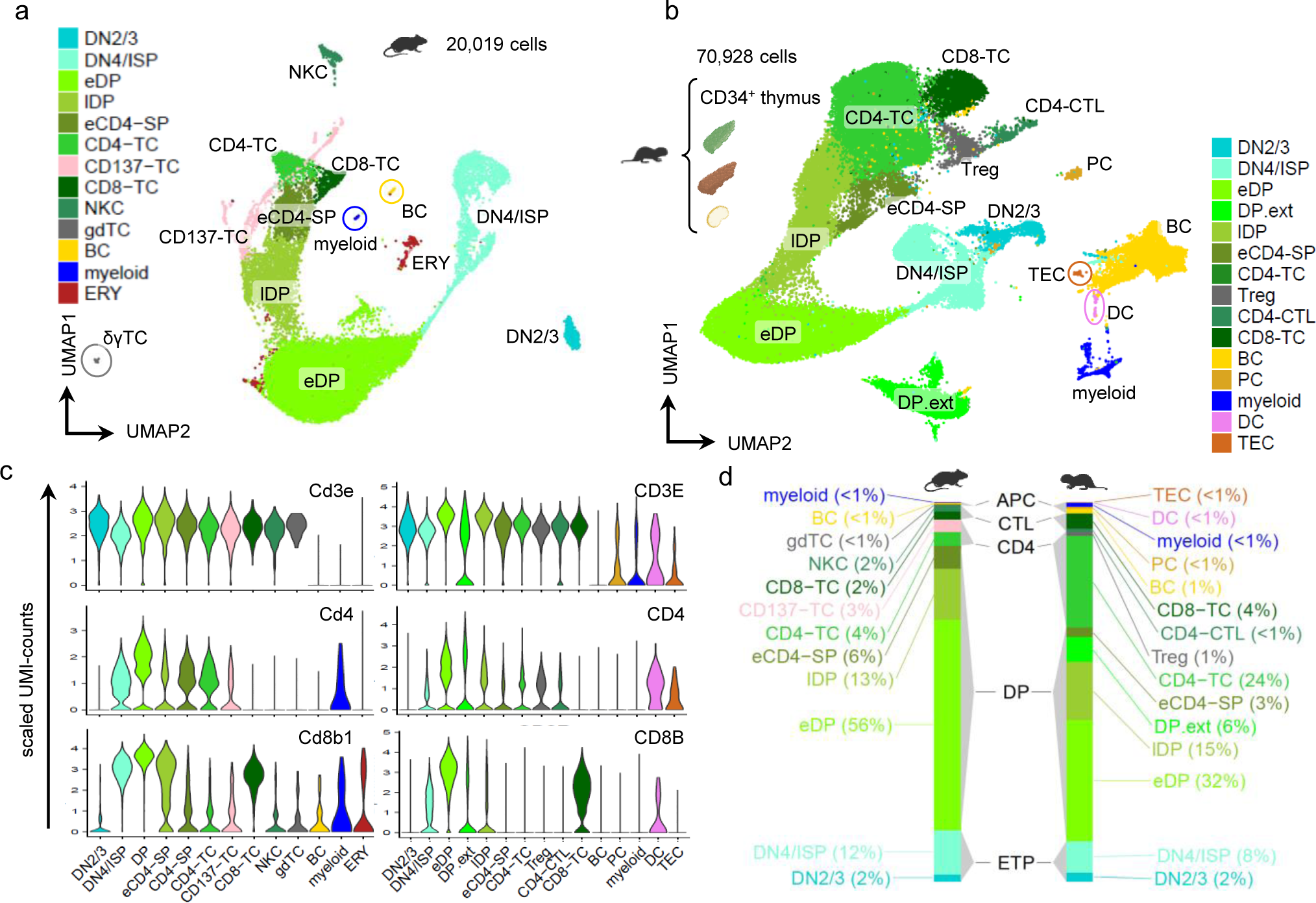
Conserved thymopoiesis between mouse and naked mole-rat. **a**, UMAP of scRNA-Seq from unfractionated thymus of 3 month-old (n=2) and 12 month-old (n=2) mice. DN2/3, CD4^−^/CD8^−^/CD44^−/+^/CD25^+^; DN4/ISP, CD4^−/lo^/CD8^−/+^/CD44^−^/CD25^−^; eDP, early double-positive CD4^+^/CD8^+^; lDP, late double-positive; eCD4-SP, early CD4 single positive; CD4- TC, CD4^+^ T-lymphocyte; CD137-TC, CD4^+^/CD137^+^ T-lymphocyte; CD8-TC, CD8^+^ T-lymphocyte; NKC, Natural Killer cell; gdTC, γδT-lymphocyte; BC, B-lymphocyte; myeloid, CD4^+^ T-lymphocyte; ERY, erythroid cells. **b**, UMAP of integrated CITE-Seq from naked mole-rat thoracic (n=4) and cervical thymus (n=4), LN (n=2) and sorted CD34^+^ thymocytes (n=1). DP.ext, extrathymic/circulating DPs; Treg, Regulatory T cell; CD4-CTL, CD4^+^ cytotoxic T-lymphocyte; PC, plasma cell; DC, dendritic cell; TEC, thymic epithelial cell. **c**, RNA expression across clusters for the mouse [left] and naked mole-rat [right] CD3, CD4 and CD8B orthologs. **d**, Bar chart of average cluster frequencies across species; mouse thymus, (n=4); naked mole-rat thoracic thymus, (n=4).

Whole thymus is usually comprised of 80-90% DPs (**Figure S3b**). Louvain clustering identified several DP communities in each species, which we labelled early (eDP) and late (lDP) according to their CD4/CD8 transcript levels. In NMRs an additional DP cluster was formed, which was detected to 6% in LNs and thus named DP.ext (extrathymic). Total DP cell frequencies were 85% in mice and 64% in naked mole-rats (**Figure 2d**). Expression of Rag1, anti-apoptotic Dek and Themis, indispensable for proper positive and negative TC selection in mice (Johnson et al., 2009), was conserved in eDPs, whereas lDPs activated Tox, Helios and Gata3 transcription (**Figure S3d-e**). Strikingly, cell type distributions between the two NMR thymus types were identical (**Figure S3h**), supporting evidence was obtained from quantitation of Thy1.1^int^/CD34^+^ and TC FACS populations, showing no difference between thymus origins (**Figure 1c**). Conversely, BCs are prevalent in LNs (**Figure S3h**), and CD125^+^ BC frequencies are higher in LNs than in BM, spleen or thymi (**Figure S4a-b**), suggesting canonical B-lymphopoietic functions in peripheral LNs.

In summary, we found an additional pair of functional thymi in naked mole-rats. Early and intermittent steps of T-lineage development appear to be conserved, however, there is a stark decrease of naked mole-rat DP proportions compared to mice.

### Cryptic γδ T-lymphocytes with a killer cell signature in naked mole-rats

Commitment towards the γδTC lineage is instructed by T cell receptor (TCR) signal strength, whereby PTCRA indicates weak and TCRγδ strong signals at the DN1 stage (Munoz-Ruiz et al., 2017). We found the constant region of the TCRγ chain (TRGC2) marking marrow ETPs (Emmrich et al., 2019), hence the most primitive thymic partition across mouse and naked mole-rat showed conserved TRGC2 overexpression (**Figure 3a**). Mature murine γδTCs are enriched for TRGC2. In naked mole-rats, however, several mature subsets contain fractions of TRGC2-expressing cells. JAML has been shown to induce γδTC activation (Witherden et al., 2010), conversely the mouse thymic γδTC cluster showed highest JAML expression (**Figure S3f**). However, we did not detect a separate γδTC population in the naked mole-rat dataset, although it had 3.5-fold more cells due to the additional cervical thymi and LNs, thereby increasing clustering resolution. Intriguingly, JAML is one of the top DN cell markers in naked mole-rats and overexpressed in CD8-TCs (**Figure S3f**). A CD4^+^ cluster comprising cytotoxic T-lymphocytes (CTL), specific for GZMA and NKG7, had a cell fraction positive for TRGC2 and JAML. Interestingly, TRGC2/NKG7 co-expressing cells were also found in CD8-TCs (**Figure S4c-d**). We therefore compared the specific cluster markers of mouse γδTC with naked mole-rat CD4-CTL (**Figure 3b**), revealing a strong correlation in a shared subset of genes encompassing S100A4/10/11, RORA, IL7R and CD44 (**Supplementary Table 2**). Conclusively, putative naked mole-rat γδTCs appeared overtly as CTLs, similar to a human Vγ9Vδ2^+^/CD45RA^+^/CD27^−^ effector memory γδTC in LNs (Caccamo et al., 2005). By contrast an abundance of murine γδTC subsets with immunomodulatory functions through mainly secreting either IFN-γ of Il-17a, but no CTLs, has been described (Pang et al., 2012). We added further evidence to this by integration of thoracic thymi scRNA-Seq from mouse and naked mole-rats, encompassing 48061 cells across 10871 common genes (**Figure 3c**). Differential abundance quantitation confirmed the between-species bias in thymic cell composition, with all mature TC clusters significantly more frequent in naked mole-rats (**Figure 3d**). Strikingly, the integrated NK/CTL cluster mapped to mouse NKCs and γδTCs, which did not co-cluster in the mouse-only dataset, and cross-mapped to naked mole-rat CD4-CTLs containing the putative killer cell γδTCs (**Figure 3e-f**). Moreover, by mapping the constant TCR chain region transcripts from all TCR loci using the most recent NMR genome (Zhou et al., 2020), we performed absolute copy number qPCR in sorted PB-TCs and saw significantly less TRGC1/2 expression in naked mole-rats (**Figure S4e**). Therefore, NMRs have a cryptic γδTC population with a CTL expression signature in the thymus, resembling human effector memory γδTCs with killer cell function. Unlike their αβTC counterparts that require peripheral activation for effector cell differentiation, γδTCs can be ‘developmentally programmed’ in the thymus to generate discrete effector subsets with distinctive molecular signatures (Munoz-Ruiz et al., 2017). Our data indicates that NMR γδTCs are overtly programed towards CTLs, potentially compensating the lack of NKCs and a diminished CD8-TC subset. CD4^+^/CTLA4^+^/CD25^+^ regulatory T cells (Treg) comprised ∼4% in NMR thymus and ∼11% in lymph nodes (**Figure S3H**), compared to 1% of human PB-WBCs or 3% of mouse lymph node cells (Greer et al., 2019), an adaptation in line with an expanded CD4-compartment across all hematopoietic tissues (Emmrich et al., 2019).

**Figure 3.**
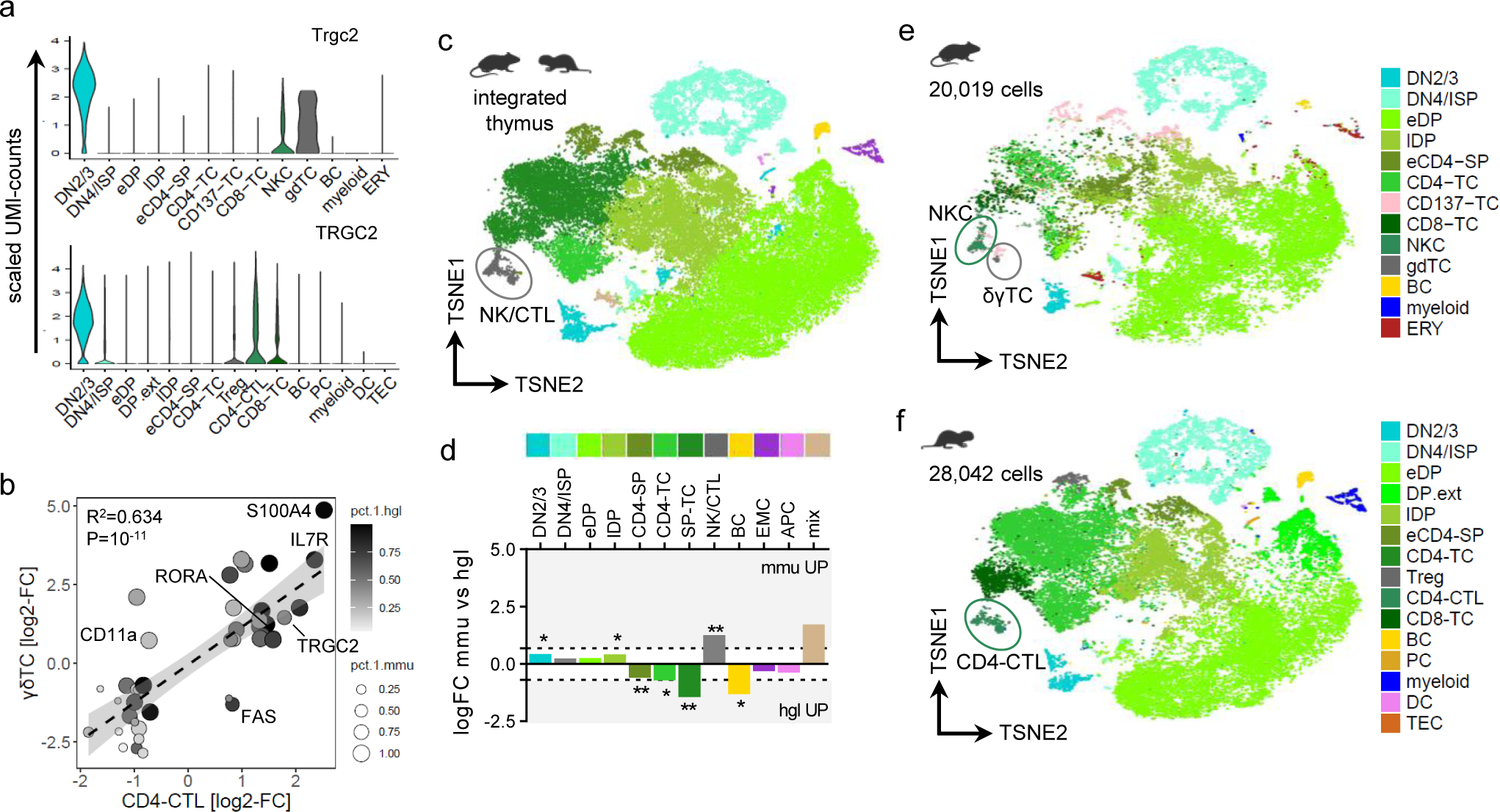
Cryptic δγT cells with killer cell signature in naked mole-rats. **a**, Expression of TRGC2 orthologs across cell types in thoracic thymi (n=4) of mouse [top] and naked mole-rat [bottom]. **b**, Correlation analysis by 45 common ortholog markers of mouse γδTCs and naked mole-rat CD4-CTLs; log2-FC, log2[fold-change] vs all other clusters in that species; fitted linear model y ∼ x. Pct.1.hgl, % of CD4-CTLs expressing naked mole-rat ortholog; pct.1.mmu, % of γδTCs expressing mouse ortholog. **c**, T-SNE of SCTransform-integrated mouse (*mmu*, n=4) and naked mole-rat (*hgl*, n=4) thoracic thymi, colorbar legend for species-integrated clusters below, encircled coordinates for NK/CTL. SP-TC, single-positice T cell; EMC, erythromyeloid cell; mix, co-clustered *mmu* erythroid with *hgl* myeloid cells and DPs. **d**, Differential cell type abundance across species; dotted lines, 2-fold change. SP-TC, NK/CTL, p<10^-3^; CD4- SP, p=0.0037; lDP, p=0.014; BC, p=0.038; CD4-TC, p=0.041; DN2/3, p=0.046. T-SNE from species-integration for the **e**, mouse or **f**, naked mole-rat partition; Cluster annotation and coloring from the single-species analysis.

### No signs of thymic immunosenescence in middle-aged naked mole-rats

In mammals, thymus tissue weight decreases with age, largely by decomposition of the thymic niche required for proper TC development, wherein thymic epithelial cells (TECs) and other stromal components are replaced by adipocytes (Palmer, 2013). Mice showed a continuous decline in thymus cellularity, which is proportional to tissue weight loss, from 3 months up to 2 years of age (19-fold, 3m-median 73`·`10^6^ vs 24m 3.9`·`10^6^ cells; **Figure 4a**), as reported earlier (Sempowski et al., 2002). To our surprise we saw a significant increase in NMR thymus cellularity between 3 and 11 years of age, for both thoracic and cervical thymi. Interestingly, cervical thymi were found to contain 10-fold more thymocytes than thoracic counterparts, which was also evident from visibly smaller thoracic lobe sizes as seen above in neonates (**Figure S4f**). However, while murine thymus volumes strongly diminished between 3 and 12 month of age, there was no shrinkage in NMR thymi in older specimen despite the 8 year age difference. We further observed cervical LNs from either species to retain size and cellularity over aging, but while LN cell numbers were consistent even between species (**Figure S4g**), NMR LNs had a much larger volume than those of mice.

**Figure 4.**
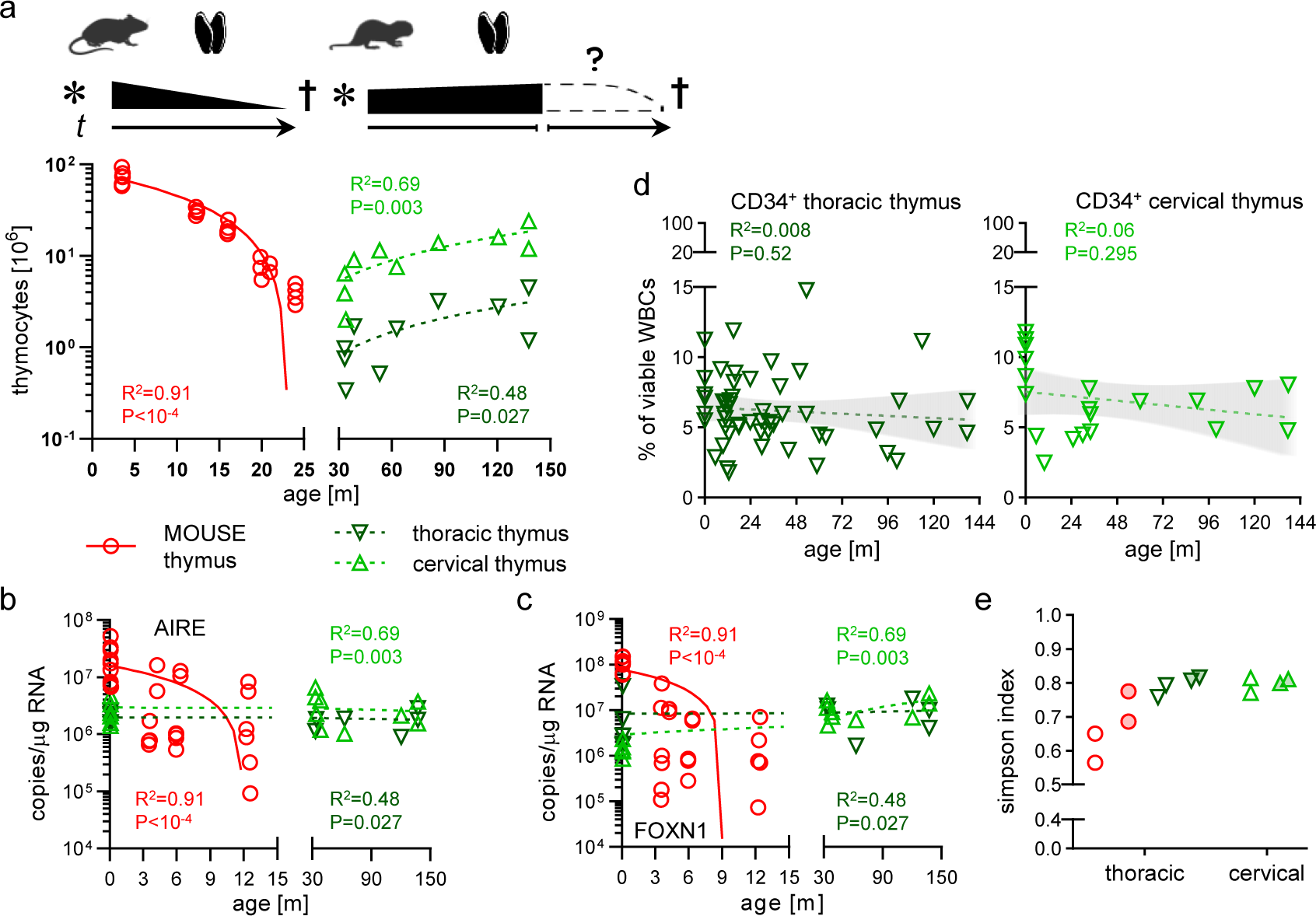
No signs of thymic immunosenescence in middle-aged naked mole-rats. **a**, Cellularity of mouse (n=24) and naked mole-rat thoracic (n=10) and cervical (n=10) thymi; R^2^ and p-value derive from linear regression. Pictograms: * birth ; †, death; *t*, time. Colors and Symbols used throughout the Figure. Absolute copy number determination for **b**, AIRE or **c**, FOXN1 ortholog mRNA in whole mouse (n=28) and naked mole-rat thoracic (n=12) and cervical (n=15) thymi. R^2^ and p-value derived from linear regression. **d**, FACS-measured CD34^+^ cell frequencies of thoracic [left, n=53] and cervical [right, n=20] thymi across age; linear regression with 95% CI as trend line; p<0.05, significance. **e**, Simpson index of cell type diversity for whole thymus single-cell transcriptomes of 3m (clear circles) and 12m (filled circles) old mice vs 3yr (clear triangles) and 11yr (filled triangles) old naked mole-rats.

We next probed several correlates of thymic aging and its concomitant loss of T-lineage potential. High autoimmune regulator gene (AIRE) activity of cortical and medullary TECs orchestrates autoantigen presentation during negative selection, and its thymic expression pattern is proportional to thymic involution (Perniola, 2018). As expected, mouse AIRE levels decreased with age as measured by absolute qPCR from whole thymi, whereas NMR AIRE was consistently expressed throughout age in naked mole-rats (**Figure 4b**). FOXN1 is the major ontogenic thymus marker by regulating TEC development and function from prenatal stage until after birth. In rodents referred to as the ‘nude locus’, genetic disruption of Foxn1 causes athymia and hairlessness in mice and rats, and a number of TC immunodeficiency syndromes have been linked to the human ortholog (Romano et al., 2013). In mice, Foxn1 levels sharply decline in the postnatal thymus, on the contrary NMR FOXN1 remains at the same expression level as detected in neonates, a clear neotenic feature (Emmrich et al., 2019) (**Figure 4c**). It is important to note that both AIRE and FOXN1 expression in NMR neonates was lower by 1-2 orders of magnitude than in mouse neonates, but whether this difference pertains to less TECs or less AIRE/FOXN1 in similar number of TECs between species warrants further investigation. We detected a TEC population in NMR thymi by scRNA-Seq with pronounced AIRE expression, whereas the mouse dataset did not feature a distinct TEC cluster (**Figure S3d-e**). Another hallmark of immunosenescence is the steady decline in frequency and functionality of DN early thymic progenitors in mice (Min et al., 2004). We showed that the most primitive NMR thymic progenitor compartment is CD34^+^, and thus quantified this fraction in thymi across an 11yr timespan. Remarkably, neither thoracic nor cervical thymi CD34^+^ populations diminished during this period (**Figure 4d**), albeit both organs showed a slight trend towards reduction, which likely becomes significant in animals aged >20yrs. The pronounced effect of thymic aging on the thymocyte pool is the dramatic decrease of naïve TCs with an accompanied accumulation of effector and memory subsets in the aged thymus. Although we did not further subset clusters due to lack of established naked mole-rat markers for memory/effector vs naïve, the Simpson index as a measure of species diversity in ecosystems can be used to score fluctuations in cell type frequencies over age (Ferrall-Fairbanks & Altrock, 2021), where represents infinite diversity and 0, no diversity. In the naked mole-rat, the Simpson index remained close to 0.8 between age groups spanning 8 years, regardless thymus type (**Figure 4e**), while in mice it rose from 0.61 to 0.73 between age groups covering 9 months. Per cell type differential abundance quantitation showed a 5-fold increase of BCs (p=0.006) and 45-fold increase of γδTCs (p=0.011) in older mice, with 5 further cell types changing >1.5-fold (**Figure S4h**). In contrast, no significant up- or down-regulated cell types were found and only 3 cell types were changed by >1.5-fold in naked mole-rats (**Figure S4i**). We further found that CD4 and CD8A mRNA levels were as consistent across age as FOXN1 and AIRE in naked mole-rat thymi (**Figure S4j-k**). These results show that in the naked mole-rat thymocyte pool composition and transcriptional patterns are maintained for over a decade, whereas in mice changes on the cellular level coincide with onset of functional thymic regress as early as within 1 year of life.

## Discussion

Here we provide the first characterization of the naked mole rat thymus. We discovered that naked mole rats have an additional pair of cervical thymi. This is an unexpected finding as mammals, including humans and mice, as a rule, have only one bilateral thymus. Cervical thymi can occasionally be detected in mice, but their frequency is rare and they have unilateral appearance (Dooley et al., 2006). Similarly, rare ectopic cervical human thymi had been reported in children (Ahsan et al., 2010). In contrast, cervical thymi are a principal component of NMR ontogenesis. Interestingly, among vertebrates, chickens have seven, sharks five, and amphibians three thymi (Boehm & Bleul, 2007). It is tempting to speculate that the presence of additional thymi in the naked mole rat may contribute to prolonged maintenance of immune function during their lifespan.

The ectopic thymus reflects a failed migration of thymic tissue from the third pharyngeal pouch endoderm during organogenesis, which can be found at any level of the pathway of normal thymic descent, from the angle of the mandible to the superior mediastinum (Saggese et al., 2002). It is possible that the naked mole-rat thymic anlage splits during migration and one remains in the throat. Alternatively, their parathyroid glands may have been repurposed to cervical thymi, which is less likely due to presence of a conserved PTH ortholog in naked mole-rat genome assemblies. Another explanation could be alterations as seen in ephrinB2 mutants (Gordon & Manley, 2011), wherein the thymus remains in the anterior pharyngeal region.

We provide evidence for a delay of thymic involution in naked mole-rats beyond the 1^st^ decade of their lifespan. Age-associated marker expression and thymic cell composition remained at the level of neonates. The absence of thymic involution up to midlife is unprecedented in mammals. This would translate into similar or even slightly heightened thymic weights and cell counts for humans in their 30’s. Thymic involution decreases output of naïve T cells and reduces the ability to mount protective responses against new antigens. In naked mole-rats we did not see thymic involution in animals >10 years old, while markers for thymic function and development, AIRE and FOXN1, were maintained at neonatal levels. Furthermore, the reduction of ETPs accompanying age-related lymphoid decline did not manifest in naked mole-rats, arguing that their intrinsic myeloid bias in the marrow does not predispose HSPCs towards less lymphoid commitment (Emmrich et al., 2019). However, naked mole-rats are not immortal and do show frailty in old age (Edrey et al., 2011). Therefore, an eventual decline in thymic cellularity and immune function is to be anticipated, albeit delayed as opposed to the lifelong steady decline in humans and mice.

Neoteny refers to retention of juvenile phenotypes in adult organisms, hence considering humans neotenic apes (Bufill et al., 2011). NMRs feature an array of neotenic traits (Skulachev et al., 2017), including aspects of their hematopoietic system (Emmrich et al., 2019). Here we found developmental FOXN1 and age-associated AIRE mRNA levels with little to no changes between neonate and 11 year-old adult animals. Similarly, thymic ETPs remain at neonate frequencies in NMRs. Thymic involution occurs in almost all vertebrates (Shanley et al., 2009), hence neotenic retention of a juvenile thymus in mature, aged animals represents a likely function of longevity by maintenance of youthful TC-mediated immune function during adulthood.

## Experimental Procedures

### Animals

Ethical and legal approval was obtained prior to the start of the study by the University of Rochester Committee on Animal Resources (UCAR). All animal experiments were approved and performed in accordance with guidelines instructed by UCAR with protocol numbers 2009-054 (naked mole rat) and 2017-033 (mouse). Naked mole rats were from the University of Rochester colonies, housing conditions as described (Ke et al., 2014). C57BL/6 mice were obtained from NIA.

### Primary cell isolation

Marrow from mice and naked mole-rats was extracted from femora, tibiae, humeri, iliaci and vertebrae by crushing. Thymus and lymph nodes were minced over a 70µm strainer and resuspended in FACS buffer. Blood from mice was drawn via retroorbital capillary bleeding, naked mole-rat blood was obtained via heart puncture.

### Histology

Imaging and analysis was performed using a using a Nikon Eclipse Ti-S microscope. Coverslips were applied with DEPEX Mounting media (Electron Microscopy Sciences), except for Alkaline Phosphatase staining where Vectashield Hard Set Mounting Medium for Fluorescence (Vector) was applied. Soft tissues were stored in 10% neutral buffered formalin, processing was done using a Sakura Tissue-Tek VIP 6 automated histoprocessor, paraffin embedding was done using a Sakura Tissue-Tek TEC 5 paraffin embedding center. A Microm HM315 microtome was used to section tissues at a thickness of 5µm, which then were floated onto a slide with a water bath at a temperature between 45°C and 55°C. Sections were deparaffinized and rehydrated to distilled water through xylene and graded ethanol (100% to 70%).

*Hematoxylin & Eosin*: Sections were stained with Mayers Hematoxylin (Sigma) for 1min and washed with tap water to remove excess blue coloring. Soft tissue sections were further decolorized with 3 dips in 0.5% acid alcohol and washed in distilled water. The nuclei of sections were blued in 1X PBS for 1 minute and washed again in distilled water. An Alcoholic-Eosin counterstain was applied for 30sec before slides were immediately dehydrated and cleared through 3 changes of 95% ethanol, 2 changes of 100% ethanol, and three changes of Xylene for 1min each.

*Cytokeratin*: Paraffin sections (4µm thick) of FFPE thymus and lymph node tissues (NMR, mouse, and human control) were stained for cytokeratin (AE1/AE3, Dako GA05361-2) on a Dako Omnis autostainer with pressure cooker antigen retrieval (TrisEDTA; pH 9). A section of normal human thymus resected for routine clinical care at the university of Rochester medical center was stained for H&E and cytokeratin for morphological comparison. The human, mouse, and NMR samples were stained in the same run on the same machine.

### Flow Cytometry

<u>Flow cytometry analysis</u> was performed at the URMC Flow Core on a LSR II or LSRFortessa (both BD), or on our labs CytoFlex S (Beckman Coulter). Kaluza 2.1 (Beckman Coulter) was used for data analysis. Staining and measurement were done using standard protocols. Red blood cell lysis was done by resuspending marrow pellets in 4ml, spleen pellets in 1ml and up to 500µl blood in 20ml of RBC lysis buffer, prepared by dissolving 4.1g NH4Cl and 0.5g KHCO3 into 500ml double-distilled H2O and adding 200µl 0.5M EDTA. Marrow and spleen were incubated for 2min on ice, blood was lysed for 30min at room temperature. Cells were resuspended in FACS buffer (DPBS, 2mM EDTA, 2% FBS [Gibco]) at 1x10^7^ cells/ml, antibodies were added at 1µl/10^7^ cells, vortex-mixed and incubated for 30min at 4°C in the dark. DAPI (Thermo Fisher) @ 1µg/ml was used as viability stain. The primary gating path for all unfixed samples was: scatter-gated WBC (FSC-A vs SSC-A) => singlets1 (SSC-W vs SSC-H) => singlets2 (FSC-W vs FSC-H) => viable cells (SSC vs DAPI) == proceed with specific markers/probes. Compensation was performed using fluorescence minus one (FMO) controls for each described panel. For antibody validation we incubated 1mio cells in 100µl Cell Staining Buffer (BioLegend; Cat# 420201) and added 5µl Human TrueStain FcX™ and 0.5µl TruStain FcX™ PLUS, followed by incubation for 10min at 4°C. We then proceeded withfluorescent antibody staining as above.

<u>Immunophenotyping</u> of naked mole-rat BM, spleen, thymus, PB and lymph nodes: CD90 FITC; CD125 PE; Thy1.1 PE-Cy7; CD34 APC, CD11b APC-Cy7. Quantification of murine BM SLAM HSCs was performed using mouse LIN Pacific Blue; Sca-1 BUV395; CD150 PE; Kit PE- Cy7; CD48 APC-Cy7. Quantification of human BM LT-HSCs was performed using human LIN Pacific Blue; CD34 APC; CD38 APC-Cy7; CD45RA FITC; CD90 PE-Cy7. Fluorescence minus one (FMO) controls were applied for fluorescent spillover compensations for each species and tissue used.

<u>Sorting</u> was performed at the URMC Flow Core on a FACSAria (BD) using a 85μm nozzle, staining was done as described. Human HSCs were sorted for population RNA-Seq as LIN^−^ /CD34^+^/CD38^Lo^/CD45RA^−^/CD90^Dim^ (Figure S5A). Naked mole-rat HSPC populations were sorted as described with a lineage cocktail comprised of CD11b, CD18, CD90 and CD125 (NMR LIN). Naked mole-rat marrow and spleen sorting panel was: NMR LIN Pacific Blue; Thy1.1 PE-Cy7; CD34 APC. Naked mole-rat blood sorting panel was: Thy1.1 PE-Cy7; CD11b APC-Cy7.

### Quantitative PCR

Mouse and Naked mole-rat sorted TCs and thymic tissue were used for RNA extraction by Trizol (Thermo Fisher). RNA was quantified using a NanoDrop One (Thermo Fisher), and 100ng was used as input for the High Capacity cDNA Reverse Transcription Kit (Thermo Fisher). RT reaction was performed according to instructions and the 20µl reaction diluted to 200µl, of which 5µl were used per qPCR reaction. We used iTaq Universal SYBR Green Supermix (Bio-Rad) on a CFX Connect® RealTime System (Bio-Rad) with a three-step cycling of 10sec 95°C, 20sec 60°C, 30sec 72°C for 40 cycles. All primers (IDTDNA) were validated to amplify a single amplicon at the above PCR conditions by gel electrophoresis. Gene sequences for primer design by Primer3Plus were retrieved from ENSEMBL, with the exception of the T cell receptor C-region genes for naked mole-rat. Here we used the WBM RNA-Seq from the transcriptome assembly below to map those genes in a recently published naked mole-rat genome (Zhou et al., 2020) using Apollo software and custom scripts. For absolute copy number quantitation, qPCR amplicons were gel-purified using the QIAquick Gel Extraction Kit (Qiagen) and subcloned into the pCR2.1 plasmid using the TOPO-TA cloning Kit (Thermo Fisher). Plasmids were prepared using the QIAprep Spin Miniprep Kit (Qiagen). Sanger sequencing was performed by Genewiz using M13 forward and reverse primers. Standard curves were prepared across a 10-fold dilution range from 20ag to 20pg of plasmid DNA. All amplicon gel images, amplicon plasmids and standard curve data is available upon request.

### Single cell RNA-Seq

10,000 DAPI^−^ thymus cells from 2 mice aged 3 month (♀&♂) and 2 mice aged 12 month (♀&♂), or 2 naked mole-rats aged 3 year (♀&♂) and 3 naked mole-rats aged 11 year (♀&♂), were subjected to CITE-Seq with 10X v3 chemistry. In addition, for all naked mole-rat specimen we collected 10,000 DAPI^−^ cervical thymus cells. For 2 naked mole-rats aged 3 year (♀&♂) we collected 10,000 DAPI^−^ cells from cervical lymph nodes. Cells were processed for TotalSeq™ CITE reagents according to the <u>manufacturers instructions</u> (BioLegend), using both human and mouse Fc blocking reagents (BioLegend). Cellular suspensions were loaded on a Chromium Single-Cell Instrument (10x Genomics, Pleasanton, CA, USA) to generate single-cell Gel Bead-in-Emulsions (GEMs). Single-cell RNA-Seq libraries were prepared using Chromium Next GEM Single Cell 3′ GEM, Library & Gel Bead Kit v3.1 (10x Genomics). The beads were dissolved and cells were lysed per manufacturer’s recommendations. GEM reverse transcription (GEM-RT) was performed to produce a barcoded, full-length cDNA from poly-adenylated mRNA. After incubation, GEMs were broken and the pooled post-GEM-RT reaction mixtures were recovered and cDNA was purified with silane magnetic beads (DynaBeads MyOne Silane Beads, PN37002D, ThermoFisher Scientific). The entire purified post GEM-RT product was amplified by PCR. This amplification reaction generated sufficient material to construct a 3’ cDNA library. Enzymatic fragmentation and size selection was used to optimize the cDNA amplicon size and indexed sequencing libraries were constructed by End Repair, A-tailing, Adaptor Ligation, and PCR. Final libraries contain the P5 and P7 priming sites used in Illumina bridge amplification. In parallel, CITE-seq library amplification is performed following SPRI bead purification of CITE-seq cDNA using Q5 Hot Start HiFi Master Mix (New England Biolabs, Ipswich, MA), SI PCR primer (IDT, Coralville, IA), and indexed TruSeq Small RNA PCR primers (Illumina, San Diego, CA) as specified(Stoeckius et al., 2017). Amplified CITE-seq libraries are purified using AMPure XP (Beckman Coulter, Indianapolis, IN) beads and quantified by Qubit dsDNA assay (ThermoFisher, Waltham, MA) and Bioanalyzer HSDNA (Agilent, Santa Clara, CA) analysis. CITE-seq libraries were pooled with 10x Genomics gene expression libraries for sequencing on Illumina’s NovaSeq 6000. Barcodes were quality filtered to keep cells between 200-5,000 detected genes/cell and <20,000 counts per cell. RNA assay was log-normalized with `“scale.factor = 1e4”`, CITE assay was `“CLR”` normalized. Variable features were detected with arguments `selection.method = “vst”, nfeatures = 3000`. Canonical correlation analysis (CCA) was used to integrate libraries (Stuart et al., 2019) from either species with `FindIntegrationAnchors` with `dims = 1:50, anchor.features = 3000, reduction = “cca”`. For naked mole-rat we integrated 4 thoracic thymus, 4 cervical thymus, 2 lymph node and 1 CD34^+^ thymocyte libraries (Figure 2b). Scores for G2M and S phases were obtained using Seurat `CellCycleScoring` as described in the respective *Seurat* <u>vignette</u>. Clustering was done using *Seurat’s* `FindClusters` function with `resolution = 0.5`. Next we used the doublet detection and removal workflow as suggested in the *<u>Bioconductor OSCA</u>* <u>vignette</u>. Briefly, we run `findDoubletCluster` from the *scDblFinder* package, followed by *in silico* simulation of doublets from the single-cell expression profiles(Dahlin et al., 2018) using `computeDoubletDensity` from *BiocSingular* package, and excluded any cluster which was identified in both methods. The DEGs for each cluster were detected by `FindAllMarkers` function with arguments `test.use = “MAST”, logfc.threshold = log(2), min.pct = 0.25, return.thresh = 0.05`. CITE feature/antibody marker detection was done as described for transcript cluster markers with the exception of `test.use = “wilcox”`. The heatmap of the top overexpressed markers for each cluster in Figure S7A shows that CCA integration worked efficiently. However, cells from sorted CD34^+^ thymocytes of the integrated clusters PC, and to a lesser extent eCD4-SP, feature a marker signature of combined DN2/3 and DN4/ISP clusters. We attribute this primarily to the difference in Chromium Single Cell 3’ Reagent Kits (10X Genomics), which were v3 chemistry for all integrated lymphoid tissues, except for the sorted CD34^+^ thymocyte library captured with v2 chemistry. We performed manual curation of the cell type annotation based on canonical markers from the literature. Mapping of naked mole-rat γδTCs to the CD4-CTL cluster was done by intersecting 620 mouse γδTC markers with 129 naked mole-rat CD4-CTL markers, yielding 45 genes in both clusters, which we regressed by a linear model from the *stats* package (Figure 3b). DA testing was performed as described(Lun et al., 2017). For the CCA-integrated naked mole-rat lymphoid dataset we show cell type abundances between age groups across both thoracic and cervical thymi (Figure S4i). Regardless of DA testing across both or separate testing of either thoracic or cervical thymi, no cell type was significantly (FDR < 0.05) changed. SCTransform was used to integrate scaled, clustered and annotated mouse and naked mole-rat unfractionated thoracic thymus datasets: `SelectIntegrationFeatures` with `nfeatures = 3000`, `FindIntegrationAnchors` with `normalization.method = “SCT”`. Cell cycle scoring, clustering and marker detection was performed as described above. Simpson’s index as calculated by

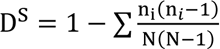 using the N(N−1) *vegan* package was determined as diversity of cell types across libraries (Figure 4e).

### Quantification and Statistical Analysis

Data are presented as the mean ± SD. Statistical tests performed can be found in the Figure legends. P values of less than 0.05 were considered statistically significant. Statistical analyses were carried out using Prism 9 software (GraphPad) unless otherwise stated.

## Acknowledgments

Many thanks to Cameron Baker for cellranger preprocessing of scRNA-Seq data, Michelle Zanche, Jeffrey Malik and John Ashton for Genomic Research Core support. The authors thank the Center for Integrated Research Computing (CIRC) at the University of Rochester for providing computational resources and technical support and the URMC Flow Core for assistance with sorting and Seahorse. This work was supported by the US National Institutes of Health grants to V.G. and A.S. and V.N.G. S.E. is a fellow of HFSP.

## Author Contributions

S.E. designed and supervised research, performed most experiments and analyzed data; F.T.Z. performed histology quantifications, animal perfusions and data analysis; A.T. contributed to bioinformatics analyses; X.Z. improved genome assembly; Q.Z. mapped TCR genes; M.D.G. performed histology and provided human BM specimen; E.M.I., Z.Z. and V.N.G. contributed to data analysis; A.S. and V.G. supervised research; S.E., A.S. and V.G. wrote the manuscript with input from all authors.

## Data Availability Statement

The data that supports the findings of this study are available in the supplementary material and Supplementary Tables of this article. The single-cell RNA-Sequencing Raw Data is available at figshare, see <u>doi link</u> and reference number **###**. Further information and requests for resources and reagents should be directed to and will be replied by the Lead Contact, Stephan Emmrich.

## Supporting Information

Supplemental Information can be found within this manuscript file.

## Supplementary Figures and Legends

**Figure S1.**
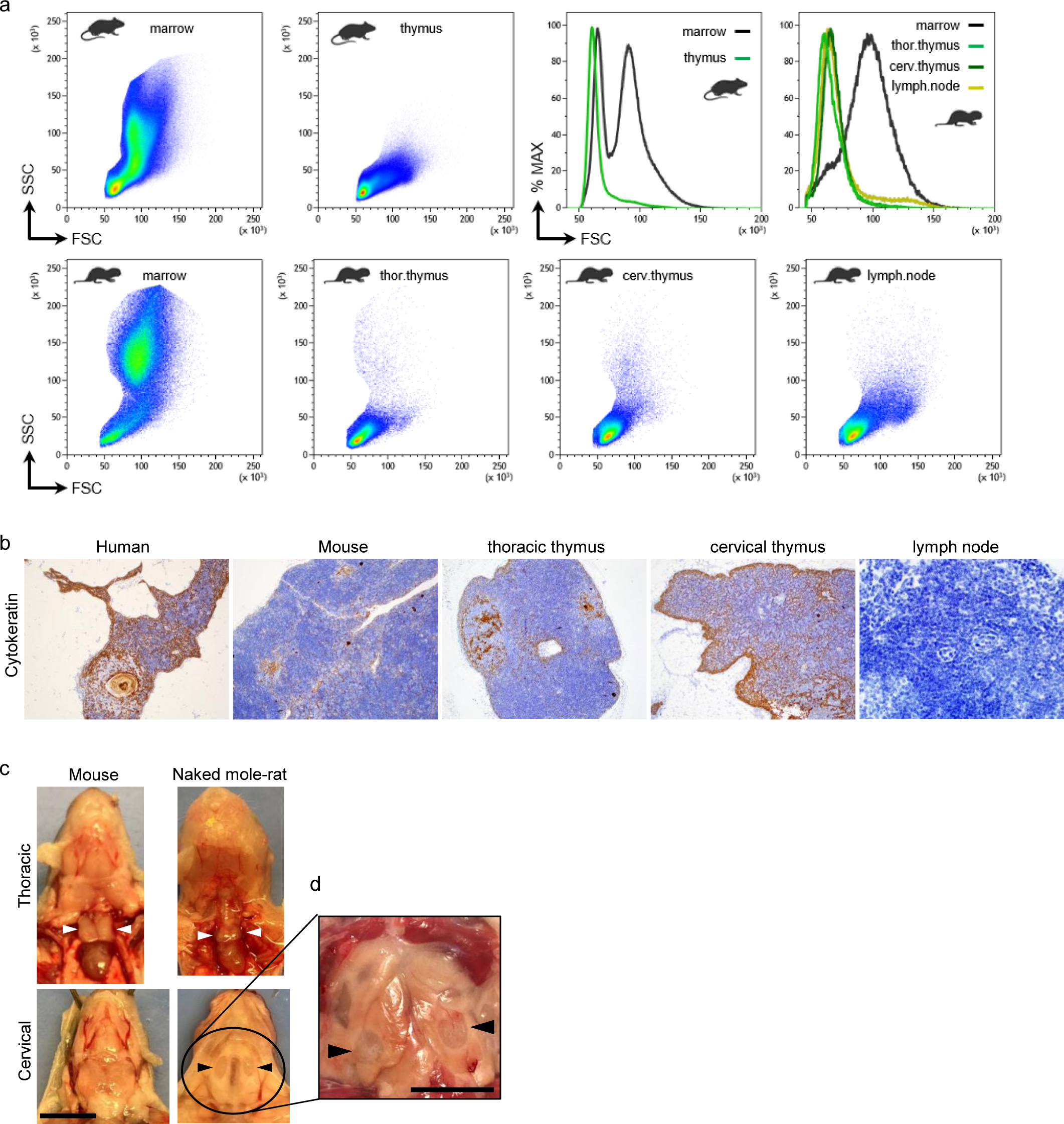
Flow cytometry and histology of mouse and naked mole-rat thymi. **a**, Viable WBCs of mouse marrow or thymus were compared for cell size [left histogram], showing mouse thymocytes with cell sizes close to marrow lymphocytes. Same can be seen for naked mole-rat thoracic or cervical thymocytes or lymph node lymphocytes, showing same size as marrow lymphocytes [right histogram]. **b**, Cytokeratin staining of thymus tissue from human [left], mouse [2^nd^ from left], naked mole-rat thoracic [middle] and cervical [2^nd^ from right], and naked mole-rat lymph node [right]; scale bar 200µm. **c**, Micrographs from the thoracic [top] or cervical [bottom] region of mouse [left] and naked mole-rat [right] neonates; scale bar 0.5cm. A bilobular thoracic thymus (white arrowheads) is present in both species, albeit drastically reduced in size in naked mole-rats. The cervical thymus (black arrowheads) is absent in mice, whereas in naked mole-rats the lobe is slightly larger on the neonate’s right side, indicating a similar transient left-right asymmetry in developmental timing and morphology as seen for murine thymic organogenesis(Gordon & Manley, 2011). **d**, Micrograph from the cervical region of an adult naked mole-rat (2.7 year-old); scale bar 0.5cm.

**Figure S2.**
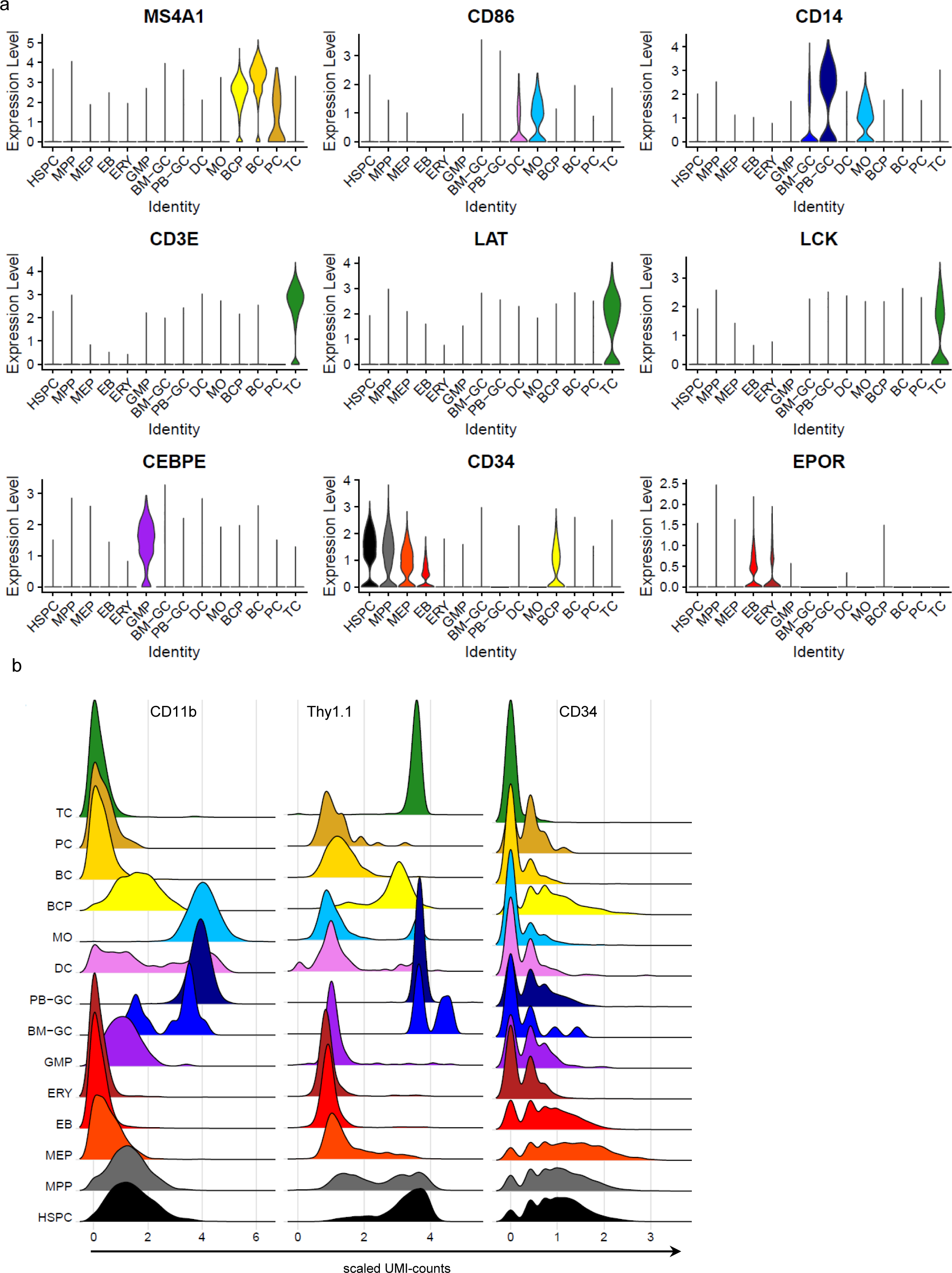
Naked mole-rat marrow and blood cell types derived from CITE-Seq. **a**, Scaled RNA-UMI counts for CD20 (MS4A1), CD86, CD14, CD3E, LAT, LCK, CEBPE, CD34 and EPOR. HSPC, hematopoietic stem and progenitor; MPP, multipotent progenitor; MEP, megakaryocytic erythroid progenitor; EB, erythroblast; ERY, erythroid cells; GMP, granulocytic monocytic progenitor; BM-GC, marrow neutrophils; PB-GC, blood neutrophils; DC, dendritic cells; MO, monocytes; BCP, B cell progenitor; BC, B cells; PC, plasma cells; TC, T cells. **b**, CITE-UMI counts for indicated cross-reactive antibodies. Naked mole-rat TCs are CITE-CD11b^−^/CD34^−^ /Thy1.1^+^. Dataset derived from (Emmrich et al., 2019).

**Figure S3.**
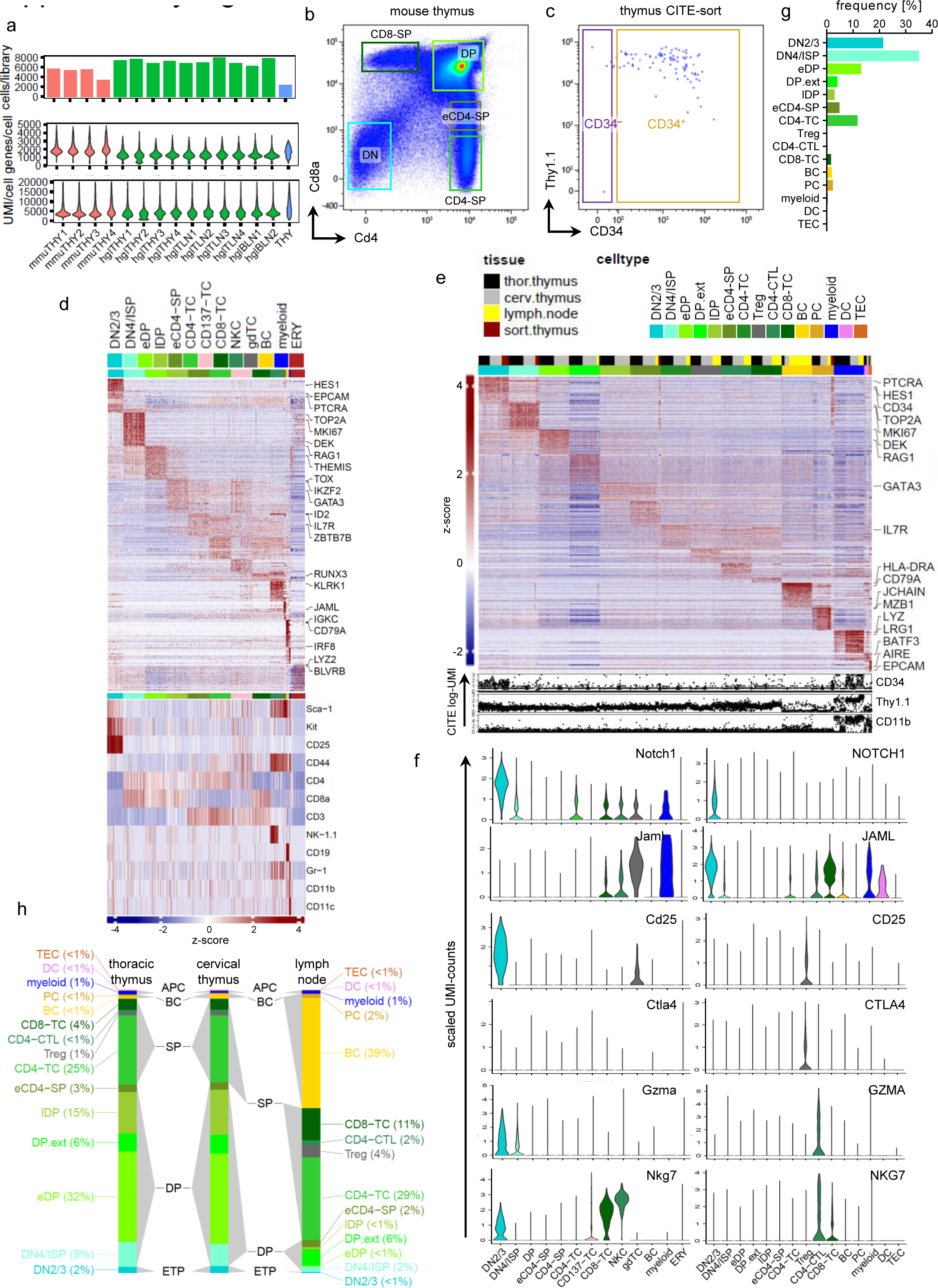
Single-cell RNA-Sequencing of mouse and naked mole-rat thymi. **a**, Quality metrics for thymus scRNA-Seq datasets from Figure 6. Top, detected genes per cell; Bottom, mRNA UMI counts per cell. THY, thoracic thymus, TLN, cervical thymus, BLN, lymph node; *mmu*, mouse; *hgl*, naked mole-rat. **b**, FACS gating of Cd4 vs Cd8a on thymocytes from a 3m old mouse. DP, Cd4^+^/Cd8a^+^ double-positive; CD8-SP, Cd4^−^/Cd8a^+^ single-positive; eCD4-SP, Cd4^+^/Cd8a^lo^ early single-positive; CD4-SP, Cd4^+^/Cd8a^−^ single-positive; Cd4^−^/Cd8a^−^ double-negative. T-lineage specification onsets with TSPs through a series of well-characterized developmental checkpoints(Schlenner & Rodewald, 2010). Termed double-negative (DN) these cells are CD4^−^/CD8^−^ and further subdivided by the expression of CD44 and CD25. Heterogeneous CD44^+^/CD25^−^ DN1 cells have the potential to form αβTC, γδTC, NKC, DCs, MAC and BC. CD44^+^/CD25^+^ DN2 cells start TCR-β, TCR-γ and TCR-δ gene segment rearrangements, DN3 cells (CD44^−^/CD25^+^) with successfully rearranged TCR-β chain enforce β-selection by association of an invariant pre-TCR-α chain (PTCRA) with CD3 signaling molecules to form the pre-TCR complex(Shah & Zuniga-Pflucker, 2014). **c**, Postsort quality control with 102 viable events of the sorted naked mole-rat CD34^+^ thymus sample, gating refers to sort decision. **d**, Top heatmap shows the top 25 cell type specific mRNA markers for mouse thymus randomly downsampled to ≤ 500 cells, canonical cell type markers from the literature are indicated. Bottom heatmap shows the cell type specific CITE features for mouse thymus randomly downsampled to ≤ 100 cells. Fold-change cut-off 2, p-value threshold 0.05. The CD137-TC cluster is partially composed of CITE-CD4^−^ cells, while TNFRSF9 (CD137) mRNA was 5.3-fold upregulated (expressed in 45.7% CD137-TC vs 1.1% in all other clusters combined; adjusted p<10^-304^). **e**, Heatmap of the top 25 cell type specific mRNA markers for the naked mole-rat lymphoid dataset from Figure 6E randomly downsampled to ≤ 500 cells, canonical cell type markers from the literature are indicated. Bottom traces depict differentially expressed CITE features as log-UMI counts. Top color bar labels tissue origin (see legend bottom right), 2^nd^ color bar labels cell type annotation. Fold-change cut-off 2, p-value threshold 0.05. **f**, Violin plots of gene expression across clusters for the mouse [left] and naked mole-rat [right] dataset. **g**, Cell frequencies of each cluster obtained from naked mole-rat sorted CD34+ thymocytes. **h**, Average cluster frequencies across unfractionated tissues; thoracic [left] and cervical [middle] thymus vs LN [right]. ETP, early T progenitor; DP, CD4^+^/CD8^+^ double positive SP, single-positive; BC, B cell; APC, antigen-presenting cell.

**Figure S4.**
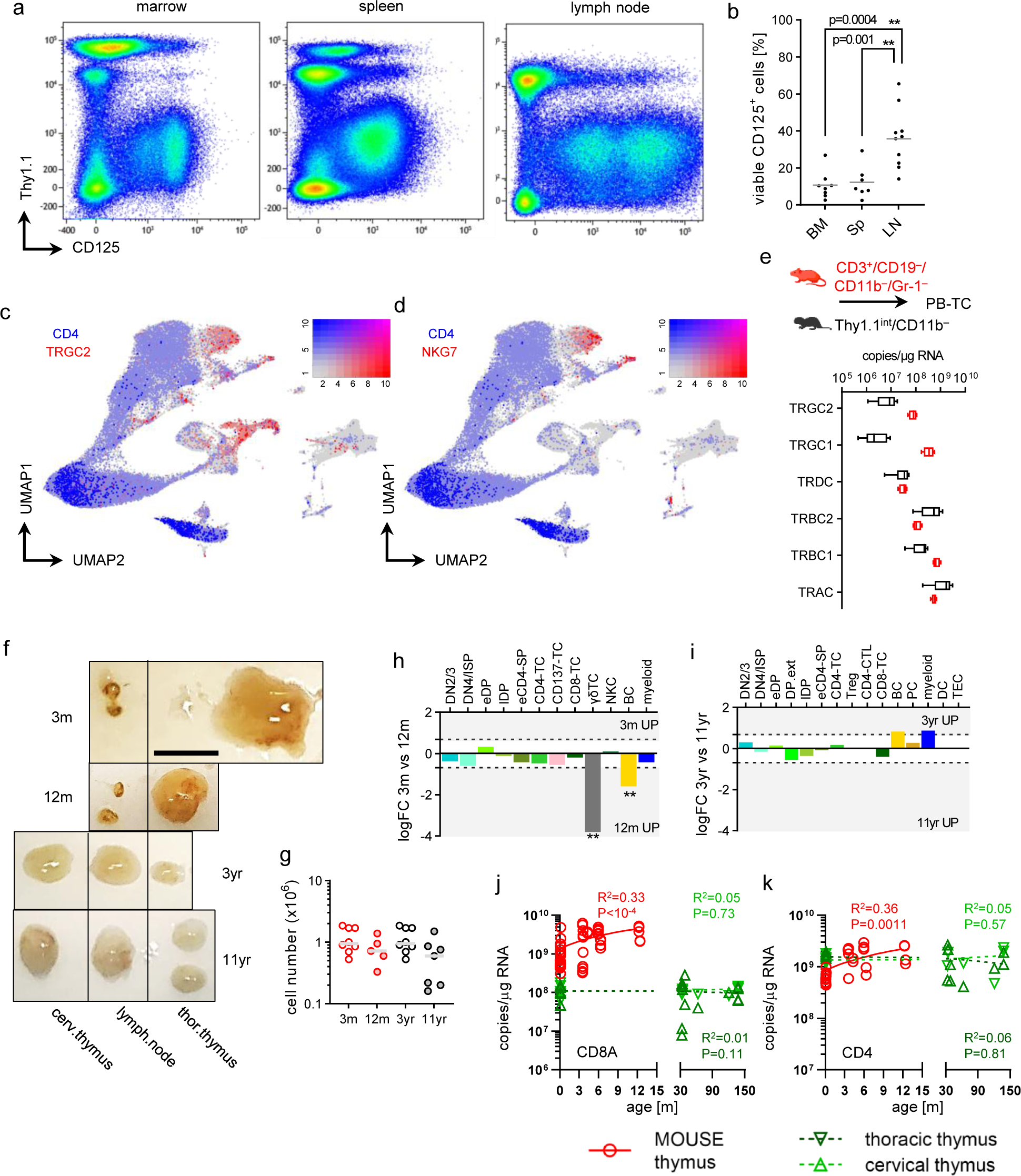
ScRNA-Seq continued and qPCR from mouse and naked mole-rat thymi. **a**, FACS gating of CD125^+^ BCs in naked mole-rat BM [left], spleen [middle] and LN [right]. Note that the lymph node CD125^+^ population has two separate CD125^dim^ and CD125^bright^ fractions. This was seen in 1/5 animals with no apparent disease condition noticed elsewise, other animal LNs showed CD125^+^ staining pattern comparable to BM. **b**, Frequencies of CD125^+^ in BM (n=8), Spleen (n=7) and lymph nodes (LN, n=10); p-value determined by Dunnett’s One-way ANOVA. **c**-**d**, UMAP-based Blendplots showing pairs conserved lineage markers; gene1 (red, high expression), gene2 (blue, high expression) and co-expressing cells (purple). See scale on the right; expression, scaled UMI counts. Cell type markers: CD4 3.7-fold up in eDP, 2.6-fold up in lDP, 2.7-fold down in CD4-CTL, 2.9-fold down in eCD4-SP and TEC, 3.1-fold down in DN4/ISP, 3.5-fold down in myeloid, 5.3-fold down in DN2/3, 5.9-fold down in PC, 6.1-fold down in CD8-TC, 6.6-fold down in BC; TRGC2 11-fold up in DN2/3, 6-fold up in CD4-CTL, 2.5-fold up in CD8-TC, 2.3-fold up in Treg; NKG7 14-fold up in CD4-CTL, 3.2-fold up in CD8-TC. **e**, Absolute qPCR of sorted T cells from mouse (n=4) and naked mole-rat (n=5) for T cell receptor constant chain orthologs; p-value derived from Tukey’s Two-way ANOVA. **f**, Micrographs of perfused thymi and lymph nodes from mice [top rows] and naked mole-rats [bottom rows]; scale bar 0.5cm. Nodes for all animals extracted from cervical/pharyngeal region. The rather puzzling finding of same-sized cervical LNs with 10-fold less cell content than cervical thymi could be explained by a one-size-fits-all encapsulation within HMW-HA, which is abundantly found across most major tissue types in naked mole-rats(Tian et al., 2013b). **g**, Cellularity of 3m (n=8) and 12m (n=5) old mice vs 3yr (n=8) and 11yr (n=7) old naked mole-rat cervical lymph nodes; p-value derived from Sidak’s One-way ANOVA. Differential cell type abundance across age for **h**, mouse or **i**, naked mole-rat single-species analysis. *mmu* BC, p=0.006; *mmu* γδTC, p=0.011. Absolute copy number determination for **j**, CD8A or **k**, CD4 ortholog mRNA in whole mouse (n=28) and naked mole-rat thoracic (n=12) and cervical (n=15) thymi. R^2^ and p-value derived from linear regression.

